# Decoupling epigenetic variation from genetic variation reveals complementary dimensions of coral eco-evolutionary dynamics

**DOI:** 10.64898/2026.08.04.742665

**Authors:** P. Buso, J. Gouspy, R. Rodolfo-Metalpa, J. de Lorgeril, V. Bonito, G. Mitta, O. Romatif, J. Pouzadoux, L. Fouré, A. Fellous, P. Auffret, C. Clerissi, E. Toulza, A. Valdivieso, J. Vidal-Dupiol, O. Rey

## Abstract

Understanding how intraspecific diversity is structured is essential for predicting the eco-evolutionary trajectories of populations, especially under rapid environmental change. While such diversity has been extensively studied from a genetic perspective, much less is known about the distribution and ecological relevance of epigenetic variation within natural populations. To address this question, we focused on two species of reef-building corals belonging to distinct functional groups, *Pocillopora acuta* and *Acropora hyacinthus*, sampled across the South Pacific (New Caledonia, Fiji, French Polynesia). Using genome-wide Enzyme-Methyl sequencing, we jointly analyzed genetic (SNPs) and DNA methylation (CpGs) variation, while explicitly disentangling genetically associated from genetically independent epigenetic variation. Genetic and epigenetic structure showed contrasting spatial patterns, reflecting distinct temporal and ecological components of population dynamics. Genetic structure was strongest between archipelagos and followed an isolation-by-distance pattern consistent with long-term evolutionary processes. In contrast, epigenetic variation converged between colonies from different archipelagos. At finer spatial scales within archipelago, genetically independent epigenetic variation exhibited stronger structure than both genetic and genetically associated epigenetic variation, likely reflecting local environmental conditions. Together, our results show that genetic and epigenetic variation provide complementary insights into the eco-evolutionary processes shaping intraspecific diversity.

## INTRODUCTION

Characterizing intraspecific diversity is fundamental for understanding eco-evolutionary dynamics and assessing the adaptive potential of populations (Eizaguirre & Baltazar-Soares, 2014; Oliver et al., 2015). While most studies have focused on genome-wide genetic diversity, epigenetic diversity in natural populations has more recently emerged as a complementary layer of intraspecific variation linking genetic background and environmental conditions, and potentially contributing to rapid adaptive responses (Bossdorf et al., 2008; Feil & Fraga, 2012; Richards et al., 2010). Variation in DNA methylation patterns, the most widely studied component of epigenetic information, arises from multiple non-mutually exclusive sources, including genetic determinism, environmental induction, and stochastic epimutations (Angers et al., 2020). However, its distribution within and among natural populations remains poorly understood, and studies have reported contrasting relationships with genetic variation (Fargeot et al., 2021; Herrera et al., 2016; Richards et al., 2012; Valverde et al., 2024). These discrepancies likely reflect both taxon-specific differences in methylation levels and functions, and the fact that epigenetic variation is often treated as a single entity, thereby conflating its distinct underlying sources. Distinguishing genetically associated from environmentally or stochastically driven epigenetic variation, particularly in comparative multispecies frameworks, may provide key insights into the contemporary processes shaping population diversity (Rey et al. 2019; Gawra et al. 2023).

Here, we investigate how genetic and epigenetic diversity are structured across spatial and environmental gradients in two reef-building coral species. Scleractinian corals provide powerful systems to address these questions. As sessile and stenothermal organisms, corals are particularly vulnerable to climate change, especially to increasing frequent and intense marine heatwaves that drive mass bleaching and mortality (Hughes et al., 2017; Van Woesik et al., 2022). Epigenetic variation may therefore play a key role by enabling rapid responses to environmental variability, complementing slower genetic processes by modulating gene expression (Dixon et al., 2018; Guerrero & Bay, 2024; Matz et al., 2018; Putnam et al., 2016). Corals also reproduce both sexually and asexually (Harrison, 2011), which can influence the spatial structure of genetic, and potentially epigenetic diversity. While genetic diversity has been extensively studied in corals (Adjeroud et al., 2014; Carr et al., 2025; Fiesinger et al., 2023; Hemond & Vollmer, 2010; Oury et al., 2026), no study has yet explicitly compared spatial patterns of genetic and epigenetic diversity.

We focused on two scleractinian corals species from different functional groups: the brooding *Pocillopora acuta* (Lamarck, 1816) and the broadcast-spawning *Acropora hyacinthus* (Dana, 1846). Colonies were sampled using a hierarchical design within and across three Pacific archipelagos spanning contrasting thermal regimes: New Caledonia, characterized by high temperature variability and infrequent heat stress; French Polynesia, with lower annual temperature variability but more recurrent heat stress events; and Fiji, with intermediate conditions. Using genome-wide SNPs and DNA methylation in CG context (CpG), we jointly analyzed genetic and epigenetic variation while explicitly distinguishing the CpGs associated with genetic variation from those that are independent. By integrating these complementary layers across spatial scales, we test the hypothesis that genetic and epigenetic variation provide decoupled, yet complementary insights into the eco-evolutionary processes that shape coral diversity. We expect genetic variation to reflect long-term demographic and evolutionary processes, primarily influenced by coral reproductive strategies and larval dispersal ability, and therefore mainly detectable at broad geographical scales. In contrast, we expect epigenetic variation independent of genetic variation, to capture ecological influences on coral colonies at finer spatial scales, likely reflecting local environmental conditions.

## RESULTS

### Overall genetic and epigenetic diversity among coral species

A total of 127 colonies of each species, *P. acuta* and *A. hyacinthus*, were analyzed to quantify genetic and epigenetic diversity across multiple spatial scales, from within-archipelago sites to inter-archipelago comparisons across the South Pacific (Fig. 1).

**Figure 1:**
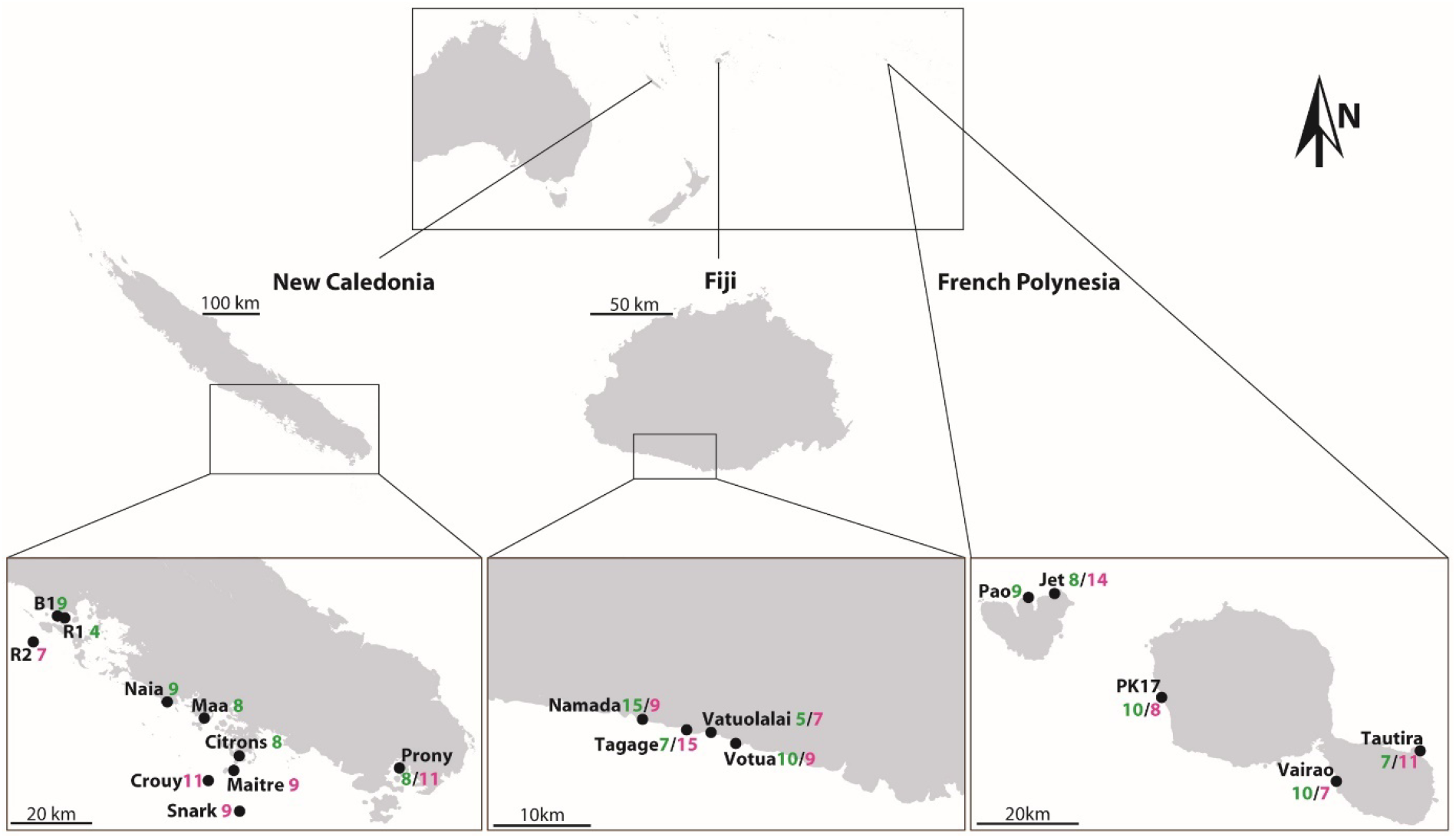
Map of sampling design across the Pacific in New Caledonia, Fiji, and French Polynesia. Insets show zoomed-in views of sampled sites within each archipelago. Black dots indicate sampling locations. Numbers associated with each site represent the number of the 127 colonies collected for each species: in green for *Pocillopora acuta* colonies and in pink for *Acropora hyacinthus*.

Based on 16,287 high-quality SNPs, relatedness analyses revealed clonality in *P. acuta*, with 81 multi-locus genotypes (MLGs) identified across archipelagos (24 MLGs/44 in French Polynesia, 36 MLGs/37 in Fiji and 21 MLGs/46 in New Caledonia (Table S1). In contrast, no evidence of clonality was detected in *A. hyacinthus* based on 12,167 SNPs. After accounting for clonality by retaining a single ramet per genet, *P. acuta* exhibited higher genetic diversity across colonies (*H*o = 0.248) than *A. hyacinthus* (*H*o = 0.153).

At the methylome level, both species exhibited DNA methylation profiles typical of invertebrates, with cytosine methylation predominantly enriched within gene bodies (Klughammer et al. 2023; Fig. 2). Overall, the two species showed broadly similar methylation patterns along metagenes, but *A. hyacinthus* consistently displayed higher methylation levels (mean = 0.223 (22%)) compared to *P. acuta* (mean = 0.07 (7%); Fig. 2A-B). MethQTL analyses detected a large fraction of CpGs that were genetically associated, particularly in *P. acuta* (106,021 out of 145,300 CpGs; *i.e*., 73.0%) compared to *A. hyacinthus* (83,337 out of 232,252 CpGs; *i.e*., 35.9%), defining two classes of CpGs hereafter referred to as “associated” and “independent” CpGs respectively.

**Figure 2:**
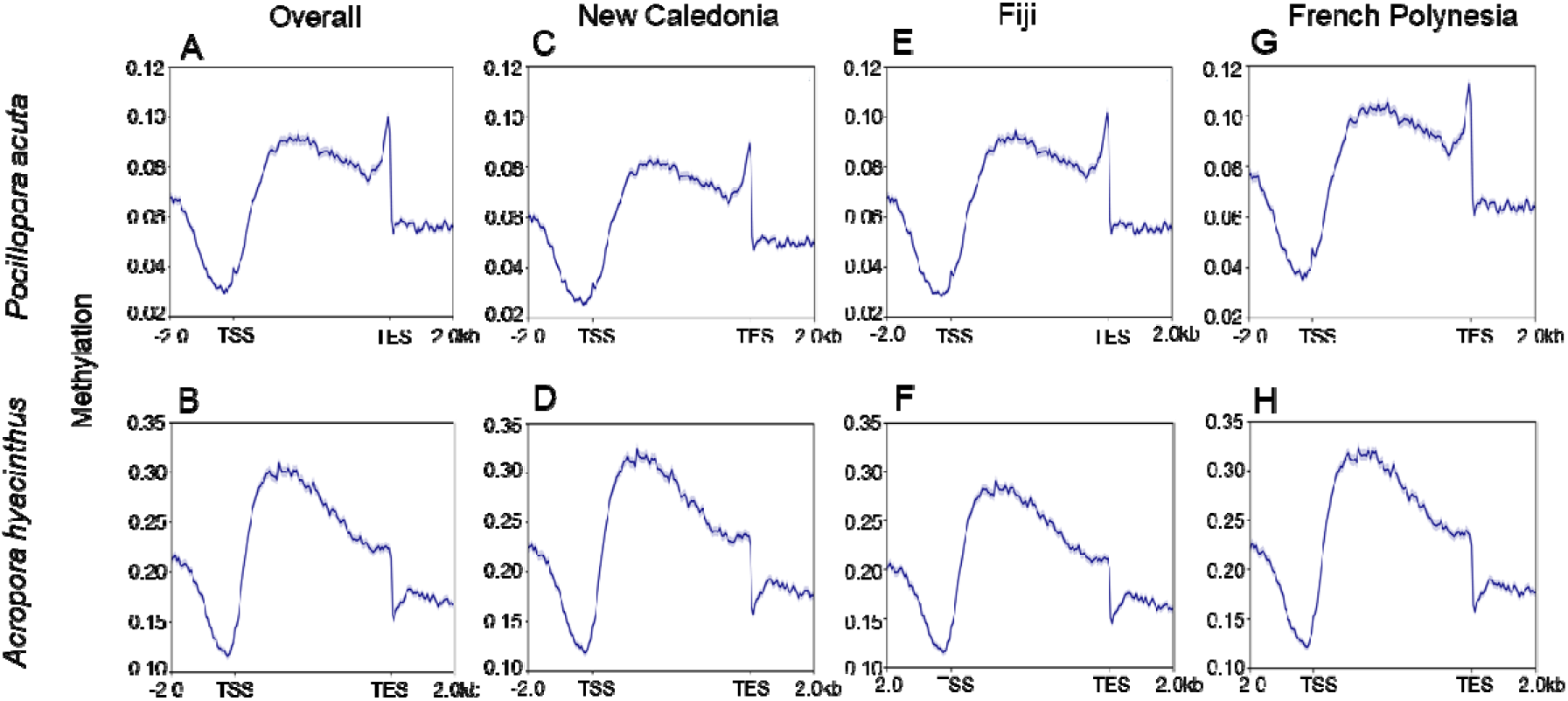
Genome-wide methylation patterns are represented across metagenes, in *P. acuta*. (A-G) and *A. hyacinthus* (B-H). Each plot depicts a 5 kb theorical gene flanked by 2 kb upstream of the transcription start site (TSS), TSS, transcription end site (TES), and +2 kb downstream. Panels A and B show all colonies combined, while panels C to H display profiles by archipelago for each species. Colonies of *A. hyacinthus* exhibits higher methylation levels at the gene body level than *P. acuta*. Gene-wide methylation rates increase along a West to East gradient from New Caledonia to French Polynesia in *P. acuta* colonies.

### Pacific-wide genetic structuring and epigenetic convergence among coral colonies

At a large spatial scale across archipelagos, genetic diversity in both species was strongly structured geographically across the South-Pacific, with distinct genetic clusters associated with each archipelago (Fig. 3A-B).

**Figure 3:**
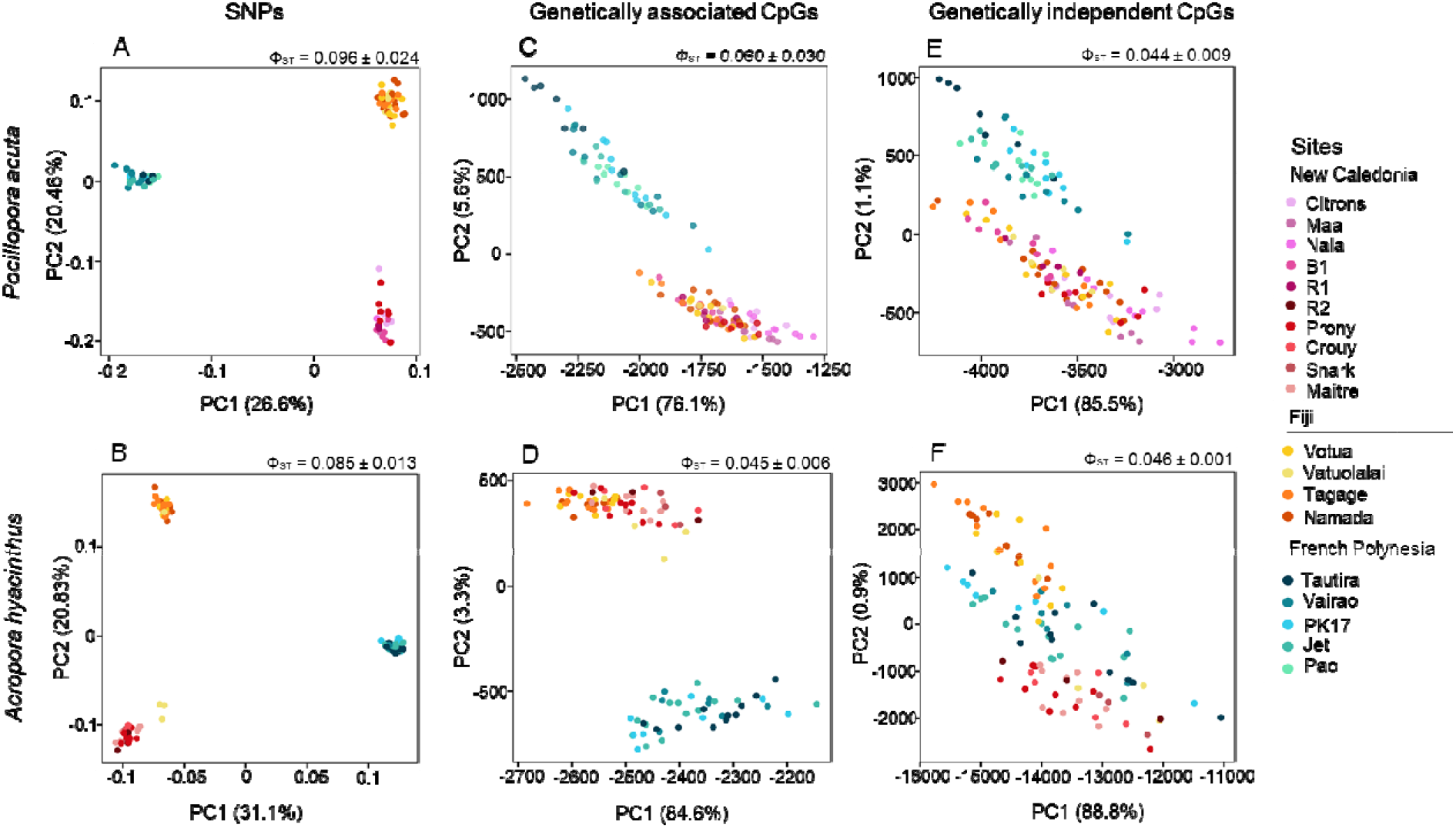
Principal component analyses (PCA) revealed that the genetic structure of colonies of *P. acuta*. (A; 16,287 SNPs) and *A. hyacinthus* (B; 12,613 SNPs) is strongly geographically structured across the Pacific. In contrast, PCAs based on associated CpG sites showed a convergence between New Caledonia and Fiji in both species (C–D), based on 106,021 CpGs in *P. acuta* and 79,277 CpGs in *A. hyacinthus*. This convergence became even more pronounced when considering only independent CpG sites (E–F), with 39,379 CpGs in *P. acuta* and 150,971 CpGs in *A. hyacinthus*.

In *P. acuta*, hierarchical AMOVAs indicated that 17.05% of the total genetic variance was attributable to differences among archipelagos (Ф_CT_ = 0.170, *p* < 0.001; Fig. 4, Table S2A). Pairwise Ф_ST_ values ranged from 0.068 between New Caledonia and Fiji to 0.115 between New Caledonia and French Polynesia, with intermediate values of 0.103 between Fiji and French Polynesia (Table S2B), consistent with isolation by distance across the Pacific (see also pairwise *F*_ST_ estimates in Table S3). Genetic diversity differed significantly among clusters, with the Fijian population (*H*_O_ = 0.269, *H*_E_ = 0.213) exhibiting the highest diversity, followed by New Caledonia (*H*_O_ = 0.252, *H*_E_ = 0.199) and French Polynesia (*H*_O_ = 0.224, *H*_E_ = 0.172). All three populations showed an excess of heterozygotes (Table S4).

**Figure 4:**
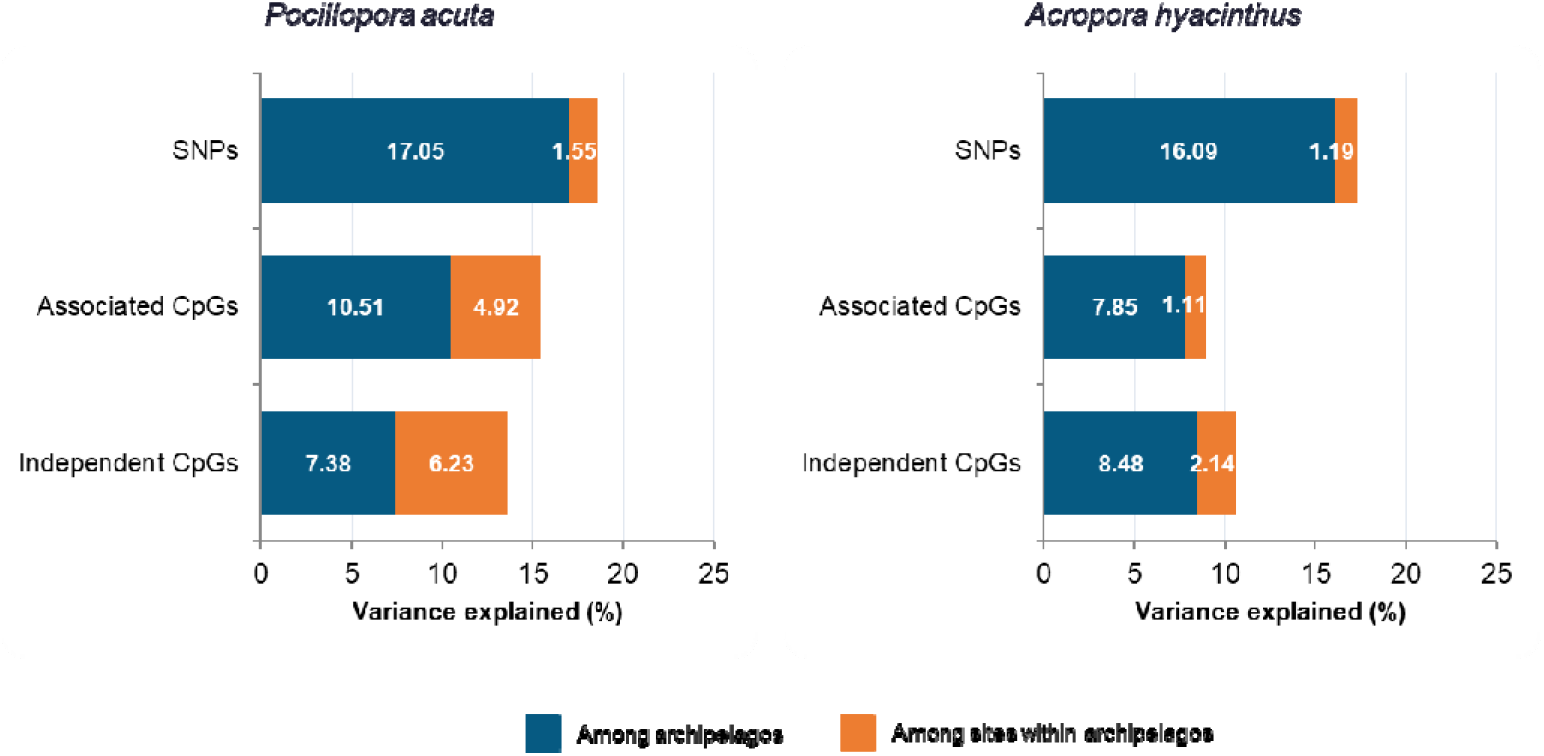
Variance partitioning using hierarchical AMOVAs computed using Euclidean distances. For each dataset (SNPs, genetically associated CpGs, and independent CpGs), in *P. acuta* and *A. hyacinthus*, molecular variance was partitioned among archipelagos and among sites within archipelagos. For each hierarchical level, the percentage of explained variance is shown on the x-axis, with datasets displayed on the y-axis. The variance components (σ²), Φ-statistics, and associated p-values are reported in Table S2. Both spatial hierarchical levels significantly contribute to genetic and epigenetic variance. The proportion of variance explained by differences among archipelagos decreases progressively from SNPs to genetically associated CpGs and further to independent CpGs. In contrast, the proportion of variance attributable to differences among sites within archipelagos is highest for independent CpGs, intermediate for genetically associated CpGs, and lowest for SNPs. These patterns suggest a partial decoupling between genetic and epigenetic hierarchical structuring.

In *A. hyacinthus*, a PCA initially revealed five genetic clusters (Fig. S1, three of which encompassed most colonies (N = 100/127), exclusive to New Caledonia (N = 29), Fiji (N = 31), and French Polynesia (N = 40). Two additional minor clusters, likely representing cryptic lineages (Rassmussen et al., 2025), were found in New Caledonia (N = 18) and Fiji (N = 9) and were excluded from subsequent analyses. A subset of the SNP and CpG matrices was therefore generated, retaining the remaining 100 colonies (Fig. 3B). The data were then filtered, and MethQTL analyses were performed on this reduced dataset (see Methods).

Hierarchical AMOVAs conducted on the three major clusters (N = 100; 12,613 SNPs) indicated that 16.09% of the total genetic variance was attributable to differences among archipelagos (Ф_CT_ = 0.161, *p* < 0.001) (Fig. 4, Table S2A). Pairwise genetic Ф_ST_ values further revealed significant differentiation between the three populations (*p* < 0.001; Table S2B), with the lowest genetic differentiation observed between New Caledonia and Fiji (Ф_ST_ = 0.070), followed by Fiji and French Polynesia (Ф_ST_ = 0.089), and finally between New Caledonia and French Polynesia (Ф_ST_ = 0.095), consistent with an isolation-by-distance pattern. Overall observed heterozygosity remained 0.141. Colonies from New Caledonia harbored the highest diversity (*H*_O_ = 0.146, *H*_E_ = 0.122), followed by those from French Polynesia (*H*_O_ = 0.142, *H*_E_ = 0.121) and lastly Fiji’s (*H*_O_ = 0.135, *H*_E_ = 0.114). All three populations exhibited a significant excess of heterozygotes (Table S4).

Epigenetic variation showed weaker structuring than genetic diversity at the South-Pacific scale and evidence of convergence among archipelagos in both species (Fig. 3 C-F). This pattern was even more pronounced for independent CpGs (Fig. 3 E-F), which consistently displayed stronger overlap and lower differentiation between archipelagos than genetic data. In *P. acuta*, archipelagos explained 10.51% of the total variance of epigenetic variation based on associated CpGs (Ф_CT_ = 0.105, *p* < 0.001) and 7.38% based on independent CpGs (Ф_CT_ = 0.074, *p* < 0.001). In *A. hyacinthus*, archipelagos explained 7.85% (Ф_CT_ = 0.079, *p* = 0.01) and 8.48% (Ф_CT_ = 0.085, *p* < 0.001) of epigenetic variation based on associated (N = 100; 79,277 CpGs) and independent CpGs (N = 100; 150,971 CpGs), respectively (Table S2A). Across both species, epigenetic differentiation remained moderate, especially based on independent CpGs (mean Φ_ST_ ≈ 0.044 and 0.046 in *P. acuta* and *A. hyacinthus*; Table S2B), supporting partial convergence of methylation profiles despite strong genetic structure. Epigenetic diversity followed a west-to-east increasing gradient in *P. acuta*, with colonies from New Caledonia displaying the lowest methylation levels across metagenes (mean = 0.061), followed by Fiji (mean = 0.068) and French Polynesia (mean = 0.078) (Fig. 2 C,E,G).

Relative to population from Fiji, colonies from New Caledonia exhibited lower methylation levels (β = −0.0076, p < 0.001), whereas colonies from French Polynesia exhibited higher methylation levels (β = 0.0099, p < 0.001) according to the linear mixed model. In *A. hyacinthus*, differences in methylation levels among archipelagos were smaller, and did not follow the same west-to-east pattern (Fig. 2 D,F,H). Colonies from French Polynesia exhibited slightly higher methylation levels than those from New Caledonia (β = −0.00172, p < 0.001) and Fiji (β = −0.0021, p < 0.001). Mean metagene methylation rates ranged from 0.210 in Fiji to 0.230 in New Caledonia and 0.231 in French Polynesia.

### Fine-scale epigenetic structuring despite limited genetic differentiation between colonies within archipelagos

Hierarchical AMOVAs also revealed significant genetic and epigenetic structure among sites within archipelagos in both species. A clear gradient of structure was observed, with the weakest differentiation in genetic data, intermediate levels in associated CpGs, and the strongest in independent CpGs.

This pattern was particularly evident in *P. acuta*, where 1.55% of genetic variation (Φ_SC_ = 0.019, p < 0.001), 4.92% of variation in associated CpGs (Φ_SC_ = 0.055, p < 0.001), and 6.23% of variation in independent CpGs (Φ_SC_ = 0.067, p < 0.001) were explained among sites within archipelagos. In *A. hyacinthus*, the corresponding values were 1.19% (Φ_SC_ = 0.014, p < 0.001), 1.11% (Φ_SC_ = 0.012, p < 0.001), and 2.14% (Φ_SC_ = 0.023, p < 0.001), respectively.

In *P. acuta,* genomic diversity (N=21/46) in New Caledonia was assessed using a subset of 7,477 variable SNPs. Based on pairwise Φ_ST_ estimates (mean 0.032 ± 0.03), only colonies from two sites (B1 and R1) were significantly genetically differentiated from a large genetic cluster encompassing colonies from all other sites (Fig. 5A; Table S5). In contrast, epigenetic variation (N = 46), whether associated (N = 47,296 CpGs) or independent (N = 59,928 CpGs) of genetic variation, was clearly structured among sites and enabled more accurate prediction of colonies’ locality of origin (Fig. 4 B-C). These patterns were supported by significant Φ_ST_ values between nearly all sites, except between Prony and Citrons, for both genetically associated CpGs (mean Φ_ST_ = 0.07 ± 0.048) and independent CpGs (mean Φ_ST_ = 0.05 ± 0.023; Table S5). Notably, sites dominated by a single clone (*e.g*., Maa, Naia) at the genetic level nevertheless exhibited substantial epigenetic diversity.

**Figure 5:**
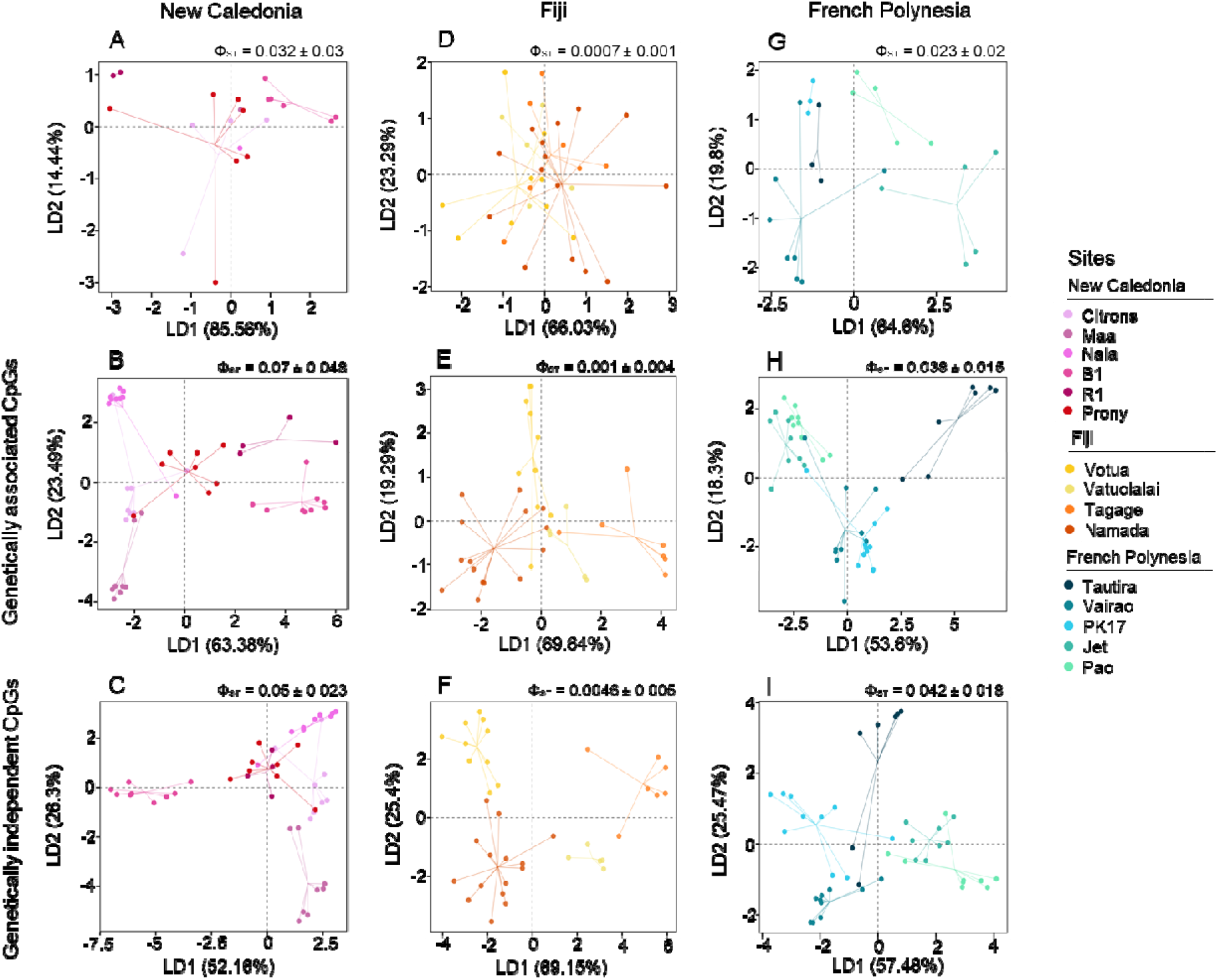
Epigenetic diversity is more structured than genetic diversity at the fine-scale level among sites within archipelagos in *P. acuta.* Each plot represents the two first discriminant axes (LD) of supervised Discriminant Analysis of Principal Components (DAPC), by archipelago in column, and for each dataset (SNPs, associated CpGs, independent CpGs) in lines.

In Fiji, the 36 *P. acuta* MLGs showed no genetic differentiation (N=9,444 SNPs) among sites (mean Φ_ST_ = 0.0007 ± 0.001). Epigenetic differentiation was not observed for associated CpGs (N = 43,336; mean Φ_ST_ = −0.001 ± 0.004) whereas independent CpGs (N = 67,988; mean Φ_ST_ = 0.0046 ± 0.005) enabled to separate significantly two of the four sites (Tagage and Namada) (Table S6). The DAPC however was able to gradually discriminate the sites with the lowest resolution in SNPs to highest resolution using independent CpGs (Fig. 5D-F).

Among *P. acuta* colonies from French Polynesia, genetic diversity (N = 24/44) was analyzed using 4,028 SNPs. Significant genetic differentiation was observed only between sites of Moorea and Tahiti Islands (mean Φ_ST_ = 0.023 ± 0.02), whereas colonies within Tahiti belonged to a single genetic cluster (Fig. 5G; Table S7). In contrast, epigenetic differentiation was higher than genetic differentiation, especially based on independent CpGs (Fig. 5H-I). Mean Φ_ST_ values were 0.038 ± 0.015 for associated CpGs (N = 47,679) and 0.042 ± 0.018 for independent CpGs (N = 63,742), all pairwise Φ_ST_ values being significant (Table S7).

In *A. hyacinthus* colonies from New Caledonia (N = 29), weaker differentiation between sites was observed at the genetic level (N = 5,187 SNPs; mean Φ_ST=_ 0.002 ± 0.002) and epigenetic level when focusing either on associated CpGs (N = 48,562 CpGs; mean Φ_ST_ = 0.01 ± 0.008) or independent CpGs (N = 158,528 CpGs; mean Φ_ST_ = 0.007 ± 0.004) (Table S8). This result was supported by the DAPC analyses (Fig. 6A-C).

**Figure 6:**
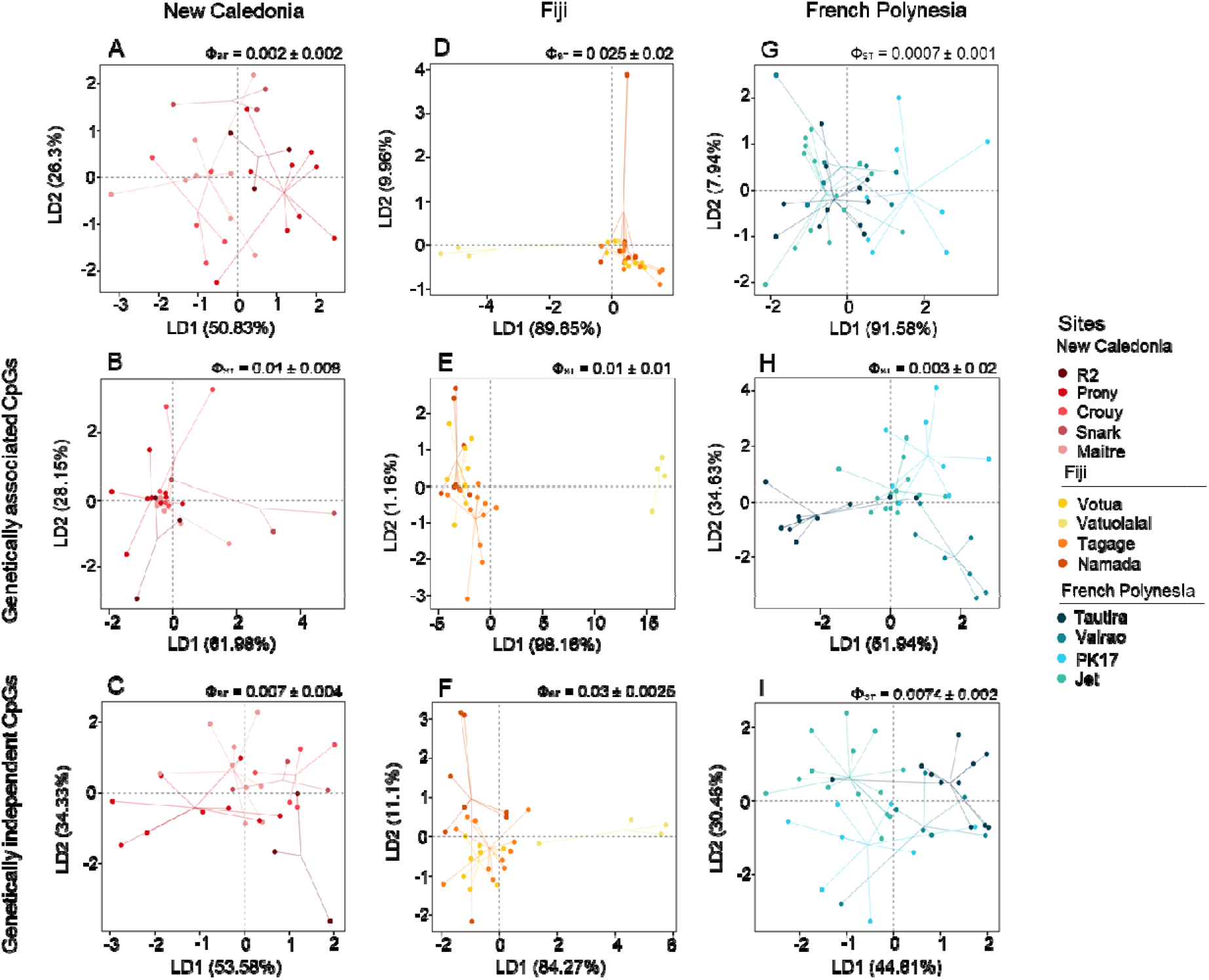
Epigenetic diversity is more structured than genetic diversity at the fine-scale level among sites within archipelagos in *A.hyacinthus.* Each plot represents the two first discriminant axes (LD) of supervised Discriminant Analysis of Principal Components (DAPC), by archipelago in column, and for each dataset (SNPs, associated CpGs, independent CpGs) in lines.

Conversely, colonies of *A. hyacinthus* in Fiji (N = 31) showed little genetic differentiation (N = 4,538 SNPs; mean Φ_ST_ = 0.025 ± 0.02) between sites. However, three of the four colonies from Vatuolalai are clearly separated along the first axis (Fig. 6D). Epigenetic diversity showed a stronger structure than genetic, particularly based on independent CpGs (N = 166,018 CpGs; mean Φ_ST_ = 0.03 ± 0.025; Fig. 6F) compared to associated CpGs (N = 44,275; mean Φ_ST_ = 0.01 ± 0.01; Fig. 6E; Table S9). Interestingly, all the four colonies from Vatuolalai appeared epigenetically differentiated from the other colonies based on both genetically associated and independent CpG datasets.

Finally, among *A. hyacinthus* colonies from French Polynesia (N = 40), structuring significantly increased from genetic (N = 7,608 SNPs; mean Φ_ST_ = 0.0007 ± 0.001) to associated CpGs (N = 57,756 CpGs; mean Φ_ST_ = 0.003 ± 0.002) and independent CpGs (N = 156,922 CpGs; mean Φ_ST_ = 0.0074 ± 0.002) (Fig. 6G-I; Table S10). Genetic differentiation was minimal, with only PK17 colonies being slightly distinct from colonies from other sites. In contrast, independent CpGs showed clear geographic structuring, with colonies distributed along a north–south gradient along the first DAPC axis and separated by exposure to dominant wind (North vs. West) along the second DAPC axis.

## DISCUSSION

Using genome-wide SNPs and DNA methylation data combined with a hierarchical sampling design, this study shows that genetic and epigenetic structure are scale-dependent and capture distinct eco-evolutionary components of intraspecific diversity. Our results illustrate that genetic variation primarily reflects long-term evolutionary processes whereas epigenetic variation is more responsive to short-term ecological processes. These patterns, consistently observed across *Pocillopora acuta* and *Acropora hyacinthus*, two coral species belonging to distinct functional groups, strengthen the robustness of this signal. The complementary contributions of genetic and epigenetic variation may therefore represent a widespread feature of intraspecific diversity, rather than a species-specific phenomenon. Together, our results support the idea that integrating multiple layers of intraspecific diversity improves our understanding of eco-evolutionary trajectories in natural coral populations across spatial scale.

As expected, genetic diversity in both coral species was strongly structured among archipelagos, following a pattern consistent with isolation by distance. Limited gene flow among archipelagos contributes to the maintenance of distinct genetic entities, as reported in several coral species (Polato et al., 2010; Voolstra et al., 2023; Oury et al., 2026; Denis et al., 2026) and other sessile or weakly dispersing organisms (Wright et al., 2015). *Pocillopora acuta* exhibited stronger population structure than *Acropora hyacinthus*, consistent with differences in reproductive strategy. Broadcast spawners (such as *A. hyacinthus*) typically show higher dispersal, promoting gene flow and reducing genetic differentiation, whereas brooders (such as *P. acuta*) often exhibit limited dispersal, high self-recruitment and sometimes clonality, leading to greater genetic structuring (Ayre & Hughes, 2000; Ayre & Miller, 2004; Adjeroud et al., 2014; Gélin et al., 2018). In *P. acuta*, genetic diversity remained homogeneous across archipelagos, whereas *A. hyacinthus* showed a lower overall genetic diversity. This cline of genetic diversity is likely associated with the thermal stress frequency that may have reduced genetic diversity through population declines and/or selective processes, primarily affecting this thermosensitive species.

Despite pronounced archipelago-driven genetic structure, both coral species consistently exhibited genetically-independent epigenetic convergence among colonies across archipelagos. This pattern is consistent with functional convergence in epigenetically mediated phenotypes, as coral species must maintain shared core biological functions essential for development (*e.g*., calcification, symbiosis, and reproduction), regardless of environmental differences among habitats within the same ecosystem. Since genetically independent CpGs reflect a mix of local environmentally-driven and stochastic dynamics (including spontaneous or labile epimutations), it rationally generates greater variation within than between archipelagos (van der Graaf et al., 2015). This interpretation is further strengthened by the conservation of methylation profiles, predominantly located within gene bodies in invertebrates, consistent with an epigenetic buffering mechanism stabilizing gene regulation under different environment despite strong genetic divergence (Dixon et al., 2018; Sarda et al., 2012). In this context, epigenetic buffering could allow populations to maintain functional integrity while genetic variation accumulates, a process that may be particularly relevant under rapid environmental change (Torda et al., 2017; Eirin-Lopez & Putnam, 2018). Genetically associated CpGs still exhibited a structure among archipelagos consistent with isolation by distance, although markedly weaker than that observed for genetic component. This likely reflects the persistence of a demographic signal embedded in genetic variation, partially blurred by weak methylation variability at genetically linked sites. The functional significance of these subtle variations at genetically associated CpG sites remains largely overlooked but clearly warrants further investigation.

In contrast, within-archipelago patterns reveal increasing epigenetic structuring as dependency on genetic variation is progressively removed. Fine-scale environmental heterogeneity likely shapes DNA methylation patterns, leading to differentiation even in the absence of genetic structure. This is consistent with theoretical and empirical studies comparing genetic and DNA methylation differentiation in wild plants and animals’ populations (Herrera et al., 2016; Rey et al., 2020; Richards et al., 2012; Sheldon et al., 2018; Thorson et al., 2017; Wogan et al., 2020). This pattern is particularly evident in *A. hyacinthus* from French Polynesia, where methylation profiles are not structured by distance but along a north–south gradient and coastline orientation. These results indicate that local ecological conditions, rather than long-term demographic processes, play a key role in shaping epigenetic variation. Notably, some epigenetic marks can be transmitted across generations, and may retain signatures of recent environmental conditions experienced by previous generations, thereby integrating ecological information over short timescales (Bossdorf et al., 2008; Fallet et al., 2020; Rey et al., 2020). Recent experimental studies on corals have shown that genetically differentiated colonies can maintain similar epigenetic marks after long-term common-garden acclimatization (Dimond & Roberts, 2020), suggesting partial stability of environmentally associated epigenetic states. Whether the observed patterns reflect within– or transgenerational plasticity (Torda et al., 2017), or recent selection on epigenetic states at the population level, remains unresolved. While such epigenetic variation could provide a substrate for natural selection or act as a temporal buffer during genetic adaptation (Anastasiadi & Piferrer, 2019; Gawra et al., 2023; Richards et al., 2010), our current data do not allow us to distinguish between these evolutionary scenarios. A spatio-temporal approach would be required to assess the stability of the epigenetic patterns and precise if epigenetic diversity contributes to coral resilience under rapid environmental change.

While the genomic localization of methylation and overall epigenetic structure were highly similar between the two species, clear quantitative differences were observed. *A. hyacinthus* exhibited higher overall methylation levels and a lower proportion of CpGs associated with genetic variation than *P. acuta*. These differences may reflect contrasting dispersal strategies (Coelho & Lasker, 2016) and differences in habitat niche breadth, resulting in species-specific levels of environmental predictability. Theoretical and empirical studies have shown that environmental predictability is tightly associated with phenotypic plasticity (Botero et al., 2015; Reed et al., 2010), itself closely linked to the capacity to generate a broader phenotypic landscape via epigenetic modulation (Leung et al. 2016; Vogt, 2022). The broader habitat niche breadth and highly dispersive larvae of *A. hyacinthus* likely expose this species to more heterogeneous and less predictable environments, potentially favoring a greater reliance on genetically independent and more plastic epigenetic mechanisms. In contrast, the narrower habitat niche breadth and high rates of self-recruitment observed in P. acuta are expected to promote a more predictable environment, potentially favoring more canalized phenotypic determinants, including both genetic variation and genetically associated epigenetic variation, with a possible long-term trajectory toward genetic assimilation (Danchin et al., 2019; Lande, 2009). Notably, the higher proportion of CpG sites independent of genetic variation in *A. hyacinthus* may also help sustain adaptive potential and population persistence despite the observed genetic depletion, as previously proposed in invasive systems (Liebl et al., 2013).

Together, our results suggest that genetic diversity at the South Pacific scale primarily reflects long-term demographic and evolutionary processes, whereas the relative lack of epigenetic structure may indicate shared functional constraints on methylation profiles across colonies sampled at a given time. At finer spatial scale, epigenetic variation provides particularly high resolution when genetic differentiation is limited. These findings highlight the value of epigenetic markers, particularly DNA methylation, for identifying ecologically distinct populations. The contrasting patterns observed between genetically associated and genetically independent epigenetic variation further show that separating these two layers is essential for interpreting the mechanisms shaping diversity. As such, integrating epigenetic information into conservation frameworks may therefore improve the design of translocation strategies, and offer promising information for monitoring natural populations complementary to genetic information.

## METHODS

### Sampling

Colonies of *P. acuta* and *A. hyacinthus* were collected in July 2022 from five sites in French Polynesia, located on the islands of Tahiti (PK17, Vairao, Tautira) and Moorea (Jet, Paopao (Pao); from eleven sites in New Caledonia on Grande Terre (Récif du Prony (Prony), Îlot Maître (Maitre), Récif Snark (Snark), Récif du Crouy (Crouy), Baie des Citrons (Citrons), Port Naia (Naia), Baie Maa (Maa), Bouraké entrée (B1), Bouraké C1 (R1), Bouraké C2 (R2); and in July 2023 from four sites in Fiji on the island of Viti Levu (Votua, Vatuolalai, Namada, and Tagage) (Fig. 1, Table S1). Colonies were sampled at a maximum depth of 3 m and with a minimum distance of 10 m between each other to minimize the collection of several ramets per genet. Each sample was obtained by cutting off a 1-3 cm fragment from the mother colony. Fragments were placed in zip-lock plastic bags fill in with surrounded sea water and bring back to the laboratory (New-Caledonia: Aquarium des Lagons; Fiji: Coral Coast Conservation Center; Tahiti-Moorea: Ifremer Pacific Center). Coral fragments were fixed on concrete supports with epoxy glue (D-D Aquascape, The Aquarium Solution) and randomly distributed in three 35 L tanks to acclimatize for 14 days at the mean temperature of the three warmer months of their archipelagos of origin (South Lagoon, New Caledonia, 28.7°C; Fiji, 28.5°C; Tahiti-Moorea, 28.7°C). This acclimatization was used to erase potential epigenetic imprinting due to seasonality and to focus only to long-term site-specific epigenetic imprinting. Water temperature was maintained using a thermal controller (T-Controller Twin 2.0, Aquamedic) coupled to a 500W titanium heater (Aquamedic). Seawater was continuously renewed at a rate of 20% of tank volume per day, and water circulation was maintained using a watermul pump (Turbelle® nanostream® 6015, Tunze) to ensure oxygen and resource availability. Temperature, salinity and pH were measured twice a day using WTW multiparameter MultiLine® 3630 IDS. At the end of the acclimatization period, samples were placed in 2 ml tubes filled with RNA later solution, stored for 24h at 4°C and then at –80°C until further processing.

### DNA extraction and sequencing

For each sample, coral fragments were blotted to remove excess of RNA later and ground for tissue-homogenization in liquid nitrogen using 50-mL stainless-steel bowls and 20-mm-diameter grinding balls at a vibrational frequency of 30 oscillations per second for 30 seconds (Retsch MM 400 mill). The resulting powder was stored in 2 mL tubes at –80°C until further processing. Ten to 20 mg of coral powder per sample was used for DNA extraction using the NucleoSpin Tissue Kit following the manufacturer’s instructions (Macherey-Nagel GmbH & Co. KG). DNA quantity and purity were checked with NanoDrop One spectrophotometer (Thermo Scientific), and quality was checked on 1% agarose gel electrophoresis before storing the DNA extracts at –20°C.

Extracted DNAs were then sent for library preparation and Enzyme Methyl-sequencing (EM-seq) at the IntegraGen plateform (Evry, France), to simultaneously access to DNA methylation and DNA sequence. EM-seq consists on enzymatic conversion of unmethylated cytosines into uracils, which are subsequently converted as thymines after PCR amplification, while methylated cytosines remain unchanged (Vaisvila et al., 2021). To assess conversion efficiency, fully unmethylated bacteriophage lambda DNA was spiked into each sample (1%) prior conversion and library construction. Libraries were sequenced using 2 × 150 bp paired-end reads on an Illumina NovaSeq platform.

### Raw data treatment and mapping

Raw reads were treated by the ACCLIMATE Nextflow pipeline (methylAtion and SNP Calling for CoraL adaptatIon to therMAl sTrEss (https://gitlab.ifremer.fr/bioinfo/workflows/acclimate). Briefly, reads quality was checked using FastQC v0.11.9 (Andrews, 2010). Sequences were trimmed to remove adaptor sequences and low-quality bases (Q < 30; bases 1–10 removed from start and end of R1 and R2) using TrimGalore v0.6.10 (Krueger, 2015). Cleaned paired-ends reads were aligned to the reference genome of each species (*P. acuta:* Stephens et al., 2022; *A. hyacinthus*: GCA_964291705.1, Welcome Sanger Institute, 2024) using Bismark v0.23.1 (Krueger & Andrews, 2011) with the following parameters: N = 0; ––score_min = L,0,-0.6. The conversion efficiency was checked by mapping the reads on the phage lambda genome (GenBank Accession NC_001416) using the same bioinformatic procedure. Finally, PCR duplicates (*i.e*., identical reads that mapped to the same regions) were removed using Bismark. Deduplicated reads (BAM files) were sorted by chromosomal position using SAMtools v1.9 (Li et al., 2009).

### Methylation and SNP calling

From the deduplicated files, methylation calling in the CG context (CpG) was performed using Bismark. Methylation coverage files of each sample were imported into R (v4.4.2, https://www.r-project.org/) and treated using “methylKit” package v1.32.1 (Akalin et al., 2012) as follows: CpG sites with a coverage < 10x and/or > 99.9th percentile were removed. Filtered methylation coverage files were then merged to form a single methylation matrix containing common CpGs shared by at least 95% of the samples, for each species independently. Coverage normalization by median was performed to account for differences in sequencing depth among colonies. The resulting methylation matrix include, for each conserved CpG site, the chromosomal coordinates and for each sample, the raw methylation rates per CpG (Beta-values).

We performed SNP calling separately for each sample using Biscuit v1.6.0 (Zhou et al., 2024), a convertion-aware variant caller. The resulting individual VCF files were filtered to retain only biallelic SNPs that passed the internal quality filters (PASS and QUAL tag of the VCF), while variants with ambiguous alternative alleles were removed. Individual VCFs were merged into a single multi-sample VCF using BCFtools v1.17 (Li, 2011). Additional filtering steps was applied using VCFtools v0.1.14 (Danecek et al., 2011) to retain SNPs with genotype quality (GQ) > 30, depth of coverage (DP) between 10x and 150x, SNP quality (QUAL) > 30, alternative allele depth > 5, and missing genotypes < 10%. SNPs with a minor allele frequency (MAF) < 0.01 were excluded, and linkage disequilibrium (LD) pruning was conducted with PLINK (v.1.9) (https://www.cog-genomics.org/plink/; Chang et al. 2015) using a sliding window of 50 SNPs, a step size of 10 SNPs, and an r² threshold of 0.3.

### Genetic diversity overall colonies between species

Based on the overall SNPs datasets, we first checked for the presence of clonality among colonies of *P. acuta* and *A.hyacinthus*. This was achieved using pairwise relatedness analysis, following the method of Manichaikul et al. (2010) as implemented in VCFtools. Kinship coefficient thresholds of > 0.354, 0.177-0.354, 0.0884-0.177, and 0.0442-0.0884 were set to detect duplicate or monozygotic twins, first-degree, second-degree, and third-degree relationships, respectively. Colonies with a kinship coefficient (Phi) > 0.354 were considered as clonal colonies (Table S1). For subsequent genetic analyses, we used the remaining colonies after clone removal by keeping one genet per ramet. Overall genetic diversity among colonies of *P. acuta* and *A. hyacinthus* at the archipelago level was assessed by calculating observed (*H*_O_) using the R package “hierfstats” v.0.5-11 (Goudet, 2005).

### Intra-specific genetic diversity and structure at the Pacific scale

Intraspecific genetic structure was explored at the Pacific scale across archipelagos using Principal Component Analysis (PCA) using PLINK (Chang et al., 2015). Hierarchical Analyses of Molecular Variance (AMOVAs, Excoffier et al. 1992) using Euclidean distance matrices were computed to partition the genetic variation among archipelagos and among sites within archipelagos using “pegas” R package v.1.3 (Paradis, 2010). Hierarchical differentiation was quantified using Φ-statistics, where Φ_CT_ represents the proportion of molecular variance attributable to differences among archipelagos relative to the total variance, and Φ_SC_ represents the proportion of variance attributable to differences among sites within archipelagos relative to the remaining variance. The significance of the variance components (the ‘archipelago’ factor and the ‘site within archipelago’ factor) were assessed using 9,999 permutations. Genetic diversity at the archipelago level was assessed by calculating observed (*H*_O_) and expected heterozygosity (*H*_E_), as well as inbreeding coefficients (*F*_IS_), using “hierfstats”. Confidence intervals (CIs) for *F*_IS_ were estimated by bootstrapping with 9,999 permutations using the *boot.ppfis* function with estimates considered significant when the CIs did not include zero. Significant differences in heterozygosity between archipelagos were tested using the Kruskal-Wallis test, followed by Dunn post hoc tests, after checking for normality and homoscedasticity of variances. Genetic differentiation between archipelagos was quantified using pairwise *F*_ST_ (Weir and Cockerham, 1984) implemented in “hierfstats” with CIs obtained by bootstrapping with 9,999 permutations using the *boot.ppfst* function. Genetic differentiation was also estimated by calculating pairwise Ф_ST_ between archipelagos based on Euclidean distances using “pegas”, for which the significance of the comparisons was assessed using 9,999 permutations.

### Intra-specific genetic diversity and structure within archipelagos

To examine intraspecific genetic diversity at finer spatial scales, analyses were repeated within each archipelago for *P. acuta* and *A. hyacinthus* colonies. Subsets of SNPs were extracted from the global VCF (after GQ, QUAL, and DP filtering) corresponding to colonies from each archipelago. These SNP subsets were further filtered to retain loci with <10% missing data, MAF > 0.01, and to minimize linkage disequilibrium, using the same parameters as in the global analyses. Discriminant Analysis of Principal Components (DAPC) was then performed under supervised frameworks, with sampling sites used as predefined populations of each archipelago using “adegenet” R package v2.1.11 (Jombart, 2008; Jombart et al., 2010). In addition, genetic differentiation among sites within each archipelago was quantified using pairwise *F*_ST_ with CIs, as well as pairwise Φ_ST_ computed on Euclidean distance matrices with 9,999 permutations.

### Genome-wide methylation patterns

Genome-wide methylation profiles were characterized using metagene analyses (*i.e*., a theoretical gene of 5,000 base pairs plus 2,000 base pairs upstream and downstream of this gene) were computed. Briefly, coverage files obtained for each sample from Bismark were converted to BedGraph files using bedtools v2.30.0 and then to bigWig files using the command bedGraphToBigWig from ucsc-bedgraphtobigwig v.472 and a species-specific chromosome size file (Quinlan & Hall, 2010). Average methylation rates from several samples were computed from bigWig and a species-specific General Feature Format (GFF) files using the command bigwigAverage under default parameters. Average methylation rates along the metagene were computed from the bigwigAverege file using the computeMatrix command (deeptools v3.5.6, Ramírez et al., 2016) and the scale-region option with the following parameters: ––regionBodyLength 5000, ––beforeRegionStartLength 2000, ––afterRegionStartLength 2000, ––skipZeros, ––missingDataAsZero, and ––binSize 50. Finally, the metagenes were plotted using the command PoltProfile from deeptools using the following parameters: ––averageType mean, ––plotType se.

### Identification of genetically associated and genetically independent CpGs

To distinguish CpG methylation variation associated with and independent from genetic variation, we used a methylation quantitative trait loci (methQTLs) approach. This approach identifies genetic variants that are statistically associated with the methylation level of one or more CpG sites. Associations were assessed using linear regression models implemented in the “GEM” R package v1.32.0 (Pan et al., 2016), following the “Gmodel”: lm(M∼G + covariate(s)), where *M* represents the methylation matrix and *G* the genotype matrix (prior to MAF and LD pruning). The models were adjusted for covariates (archipelagos for *A. hyacinthus* and archipelagos and sampling sites for *P. acuta*). Missing methylation values were imputed using median methylation rates by site within each subset. CpGs with unvariable methylation across individuals, including those with null methylation rates, were discarded. Significance thresholds for SNP-CpG associations were determined using Bonferroni correction. CpG sites showing significant genotype–methylation associations were classified as genetically associated CpGs (“associated CpGs”), whereas the remaining CpG sites were considered genetically independent CpGs (“independent CpGs”), resulting in two separate methylation matrices for downstream analyses.

### Epigenetic diversity and structure at the Pacific scale

Intraspecific epigenetic diversity was characterized using the same multivariate framework applied to genetic data, using all colonies (all ramet of each genet). First, PCA was performed on methylation matrices using the “*prcomp*” function in the base “stats*”* R package v4.4.3, separately using associated CpGs and independent CpGs matrices. Hierarchical AMOVAs with 9,999 permutations were conducted on Euclidean distance matrices to investigate how epigenetic variation is hierarchically partitioned among archipelagos and among sites within archipelagos for each species. Pairwise Ф_ST_ were computed between colonies from the three archipelagos. For these two latest analyses, beta-values were log2-transformed to M-values before calculating Euclidean distances and performing AMOVAs.

### Epigenetic diversity and structure at the archipelago scale

At finer spatial scales, analyses were repeated within each archipelago for the two coral species. Global methylation matrices were subset to obtain CpGs matrices per archipelago and filtered to remove CpGs with unvariable CpGs across all individuals. MethQTL analyses were repeated for each subset using the corresponding CpGs and SNPs matrices with the sampling site as covariate. *P*-values were adjusted independently for each subset using Bonferroni correction. DAPC were performed under supervised frameworks, with sampling sites defined as prior groupings. Epigenetic differentiation between sites was estimated using pairwise Ф_ST_ from Euclidean distances on M-values, and statistical significance was evaluated using 9,999 permutations.

## DATA AVAILABILITY

EM-seq data have been deposited under the SAVE BioProject accession XXX. EM-seq data are available under accession number XXX-XXX. ACCLIMATE Nextflow pipeline for EM-seq treatment is available here https://gitlab.ifremer.fr/bioinfo/workflows/acclimate. Code used for this article is available from https://github.com/paulinebuso/Genetic-and-epigenetic-structure-of-coral-species

### ACKNOWLEDGEMENTS

We thank Olivier Chateau and the Aquarium des Lagons for their collaboration on running ex-situ aquarium experiments. We would like to thank Pierre-Alexandre Gagnaire from the Institut des Sciences de l’Évolution de Montpellier for fruitful discussions and valuable comments on the manuscript. We thank the bioinformatics service of Ifremer (SEBIMER) for their assistance with bioinformatics analyses.

## AUTHOR CONTRIBUTION

Funding acquisition: J.V-D

Conceptualization: O.R and J.V-D

Sampling and Experimentation: J.V-D, O.R, R.R.M, J.D.L, A.F, C.C, L.F, V.B, O.R, J.P

Methodology: P.B, P.A, O.R, J.V.D, E.T, G.M

Investigation: P.B, J.G, P.A, O.R, J.V.D, E.T, A.V

Supervision: O.R. and J.V-D

Writing original draft: P.B, J.V-D, O.R

Writing review and editing: all authors.

