## Supplementary figures and tables for "Decoupling epigenetic variation from genetic variation reveals complementary dimensions of coral eco-evolutionary dynamics"

Pauline Buso *et al.*

**This PDF file includes:**

Fig. S1

Tables S1 to S13

**Other Supplementary Material for this manuscript includes the following:**

Data. S1

**Fig. S1**. PCA of genetic structure of *A.hyacinthus* (N = 127; 12,167 SNPs)

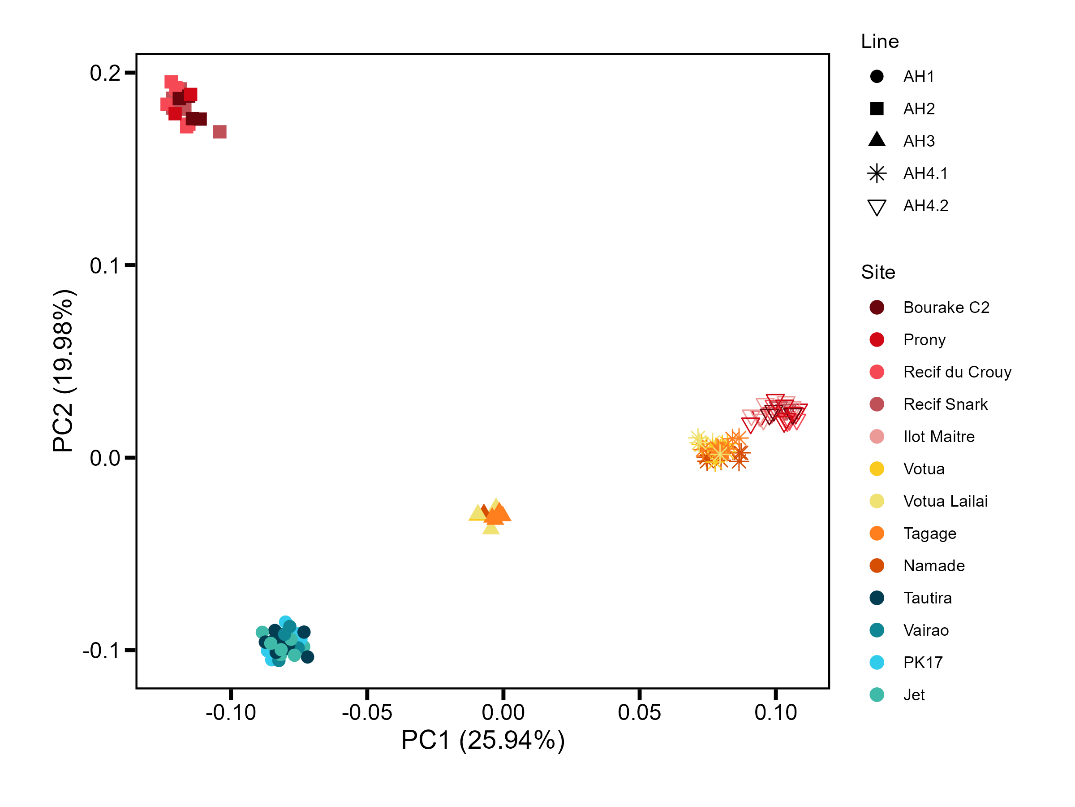

New Caledonia

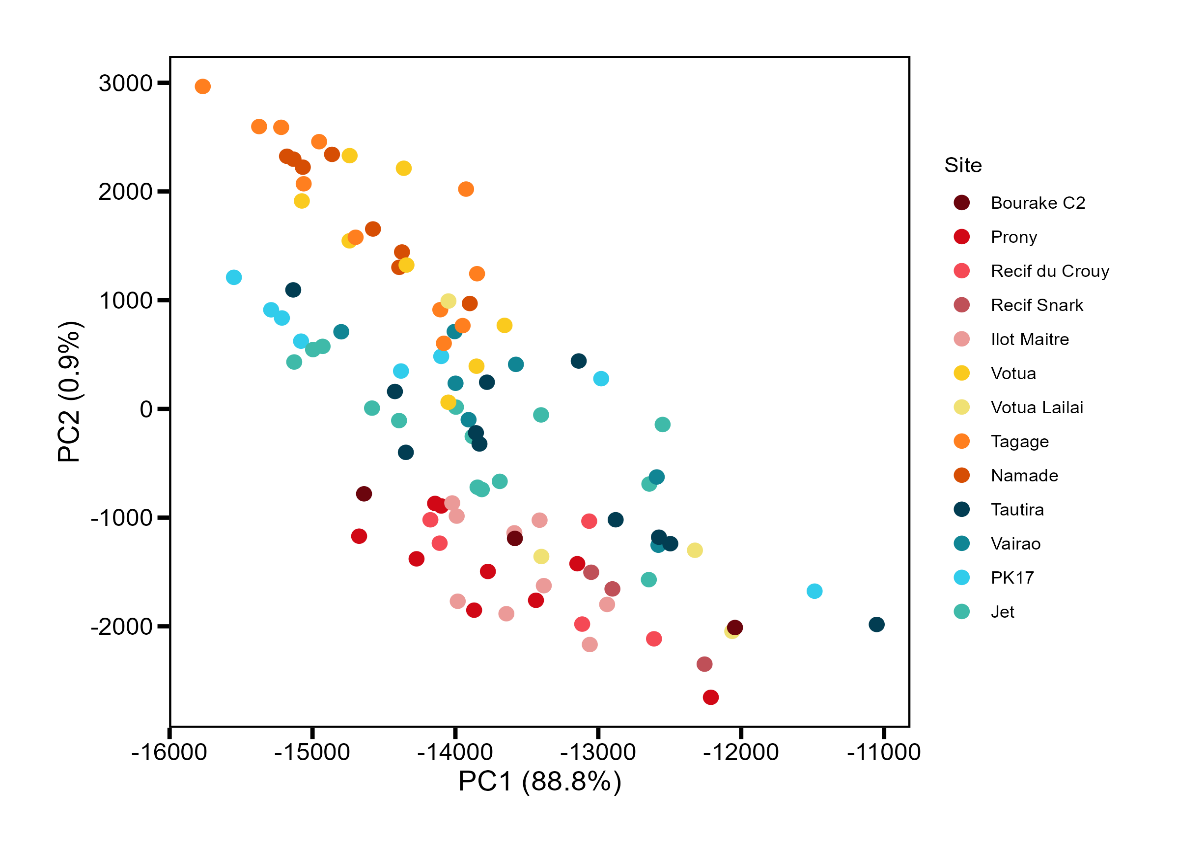

R2

Prony

Crouy

Snark

Maitre

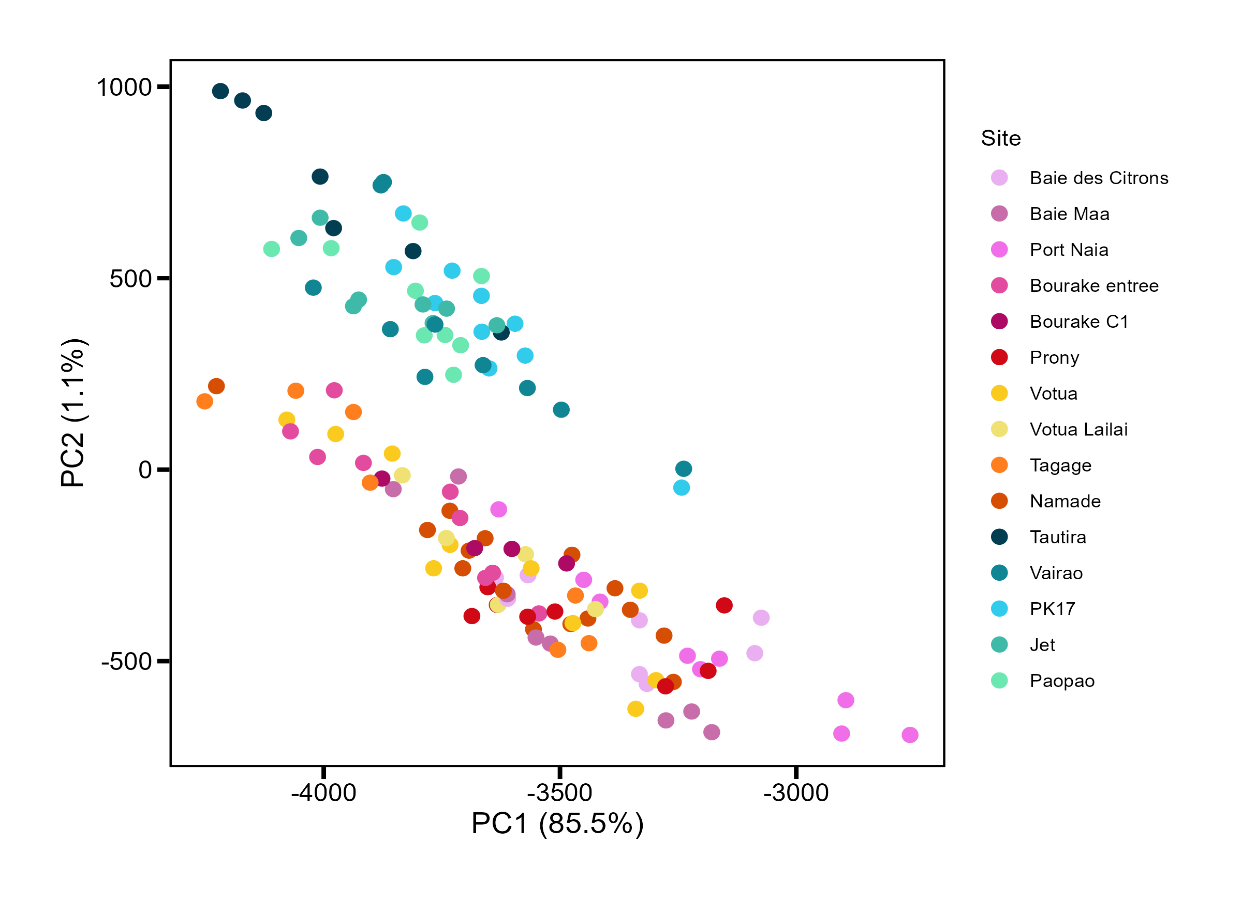

Votua

Vatuolalai

Namada

Tagage

Tautira

Vairao

PK17

Jet

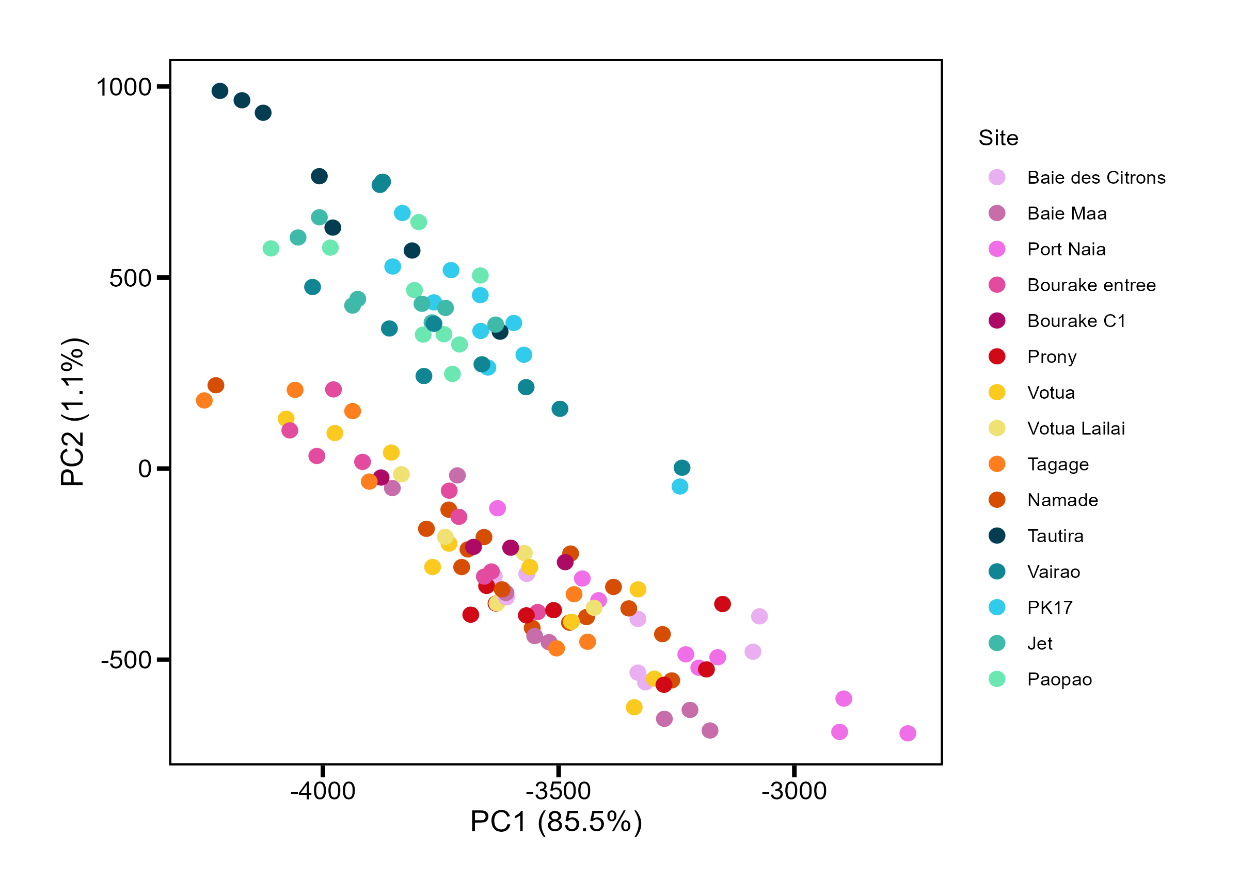

Fiji

French Polynesia

Sites

AH2

N=18

AH3

N=9

AH1

N=40

N=31

N=29

AH4

**Table S1**. Hierarchical AMOVAs performed on Euclidean distances computed from SNPs, genetically associated CpGs, and independent CpGs data of *P. acuta* and *A.hyacinthus*. Molecular variance is partitioned among archipelagos and among sites within archipelagos. The residual variance corresponds to “within sites” variability. For each hierarchical level, variance components (σ²), percentage of variance explained, Φ-statistics, and associated p-values are reported.

| Dataset | Source of variation | Variance components (σ^2^) | % Variation | Ф-statictic | *p*-value |
| --- | --- | --- | --- | --- | --- |
| *P.acuta* | | | | | |
| SNPs | Among archipelagos | 941.09 | 17.05 | 0.170 | < 0.001 |
|  | Among sites within archipelagos | 85.63 | 1.55 | 0.019 | < 0.001 |
|  | Residual | 4493.60 | 81.40 |  |  |
| Associated CpGs | Among archipelagos | 40202 | 10.51 | 0.105 | < 0.001 |
|  | Among sites within archipelagos | 18815 | 4.92 | 0.055 | < 0.001 |
|  | Residual | 323532 | 84.57 |  |  |
| Independent CpGs | Among archipelagos | 26434 | 7.38 | 0.074 | < 0.001 |
|  | Among sites within archipelagos | 22342 | 6.23 | 0.067 | < 0.001 |
|  | Residual | 309603 | 86.39 |  |  |
| *A.hyacinthus* | | | | | |
| SNPs | Among archipelagos | 455.67 | 16.09 | 0.161 | < 0.001 |
|  | Among sites within archipelagos | 33.78 | 1.19 | 0.014 | < 0.001 |
|  | Residual | 2342.52 | 82.72 |  |  |
| Associated CpGs | Among archipelagos | 35834.6 | 7.85 | 0.079 | 0.01 |
|  | Among sites within archipelagos | 5049.2 | 1.11 | 0.012 | < 0.001 |
|  | Residual | 415590.5 | 91.04 |  |  |
| Independent CpGs | Among archipelagos | 169696 | 8.48 | 0.085 | < 0.001 |
|  | Among sites within archipelagos | 42774 | 2.14 | 0.023 | < 0.001 |
|  | Residual | 1789325 | 89.39 |  |  |

**Table S2.** Pairwise Φ_ST_ between archipelagos in *Pocillopora acuta* for the three datasets (SNPs, genetically associated CpGs and genetically independent CpGs).

| SNPs (81 colonies; N=16,287); mean ΦST = 0.096 ± 0.024 (p < 0.001) | | | |  | |  |
| --- | --- | --- | --- | --- | --- | --- |
|  | New Caledonia | | | Fiji | | French Polynesia |
| New Caledonia |  | | |  | |  |
| Fiji | 0.068 | | |  | |  |
| French Polynesia | 0.115 | | | 0.103 | |  |
| Genetically associated CpGs (127 colonies; N=106,021); mean ΦST = 0.060 ± 0.030 (p < 0.001) | | | | | |  |
|  | New Caledonia | | | Fiji | | French Polynesia |
| New Caledonia |  | | |  | |  |
| Fiji | 0.025 | | |  | |  |
| French Polynesia | 0.084 | | | 0.070 | |  |
| Genetically independent CpGs (127 colonies; N=39,279); mean ΦST = 0.044 ± 0.009 (p < 0.001) | | | | | |  |
|  | New Caledonia | | | Fiji | | French Polynesia |
| New Caledonia |  | | |  | |  |
| Fiji | 0.036 | | |  | |  |
| French Polynesia | 0.053 | | | 0.043 | |  |

**Table S3.** Pairwise *F*_ST_ between archipelagos in *P.acuta* *A.hyacinthus*

| *Pocillopora acuta* (N=81; 16,287 SNPs) | | | |
| --- | --- | --- | --- |
| **Population** | **Pairwise *F*_ST_** | **CI lower** | **CI upper** |
| New Caledonia / Fiji | 0.050 | 0.049 | 0.052 |
| New Caledonia / French Polynesia | 0.088 | 0.086 | 0.090 |
| Fiji / French Polynesia | 0.075 | 0.074 | 0.077 |

**Table S4**. Heterozygosity of *P.acuta* at archipelago level. Observed heterozygosity differed significantly among *P.acuta* populations (Kruskal-Wallis test, χ² = 893.79, p < 0.001). Pairwise comparisons (Dunn-test with BH correction) revealed significant differences between all population pairs.

| *Pocillopora acuta* (N=81; 16,287 SNPs) | | | | | |
| --- | --- | --- | --- | --- | --- |
| **Population** | ***H*o** | ***H*e** | ***F*_IS_** | **CI lower** | **CI upper** |
| New Caledonia | 0.252 | 0.199 | -0.268 | -0.272 | -0.263 |
| Fiji | 0.269 | 0.213 | -0.259 | -0.263 | -0.255 |
| French Polynesia | 0.224 | 0.172 | -0.301 | -0.303 | -0.293 |

**Table S5**. Pairwise Φ_ST_ between archipelagos in *Acropora hyacinthus* for the three datasets (SNPs, genetically associated CpGs and genetically independent CpGs).

| SNPs (N=12,613); mean ΦST = 0.085 ± 0.013 (p < 0.001) | | |  |
| --- | --- | --- | --- |
|  | New Caledonia | Fiji | French Polynesia |
| New Caledonia |  |  |  |
| Fiji | 0.070 |  |  |
| French Polynesia | 0.095 | 0.089 |  |
| Genetically associated CpGs (N=79,277); mean ΦST = 0.045 ± 0.006 (p < 0.001) | | |  |
|  | New Caledonia | Fiji | French Polynesia |
| New Caledonia |  |  |  |
| Fiji | 0.038 |  |  |
| French Polynesia | 0.049 | 0.047 |  |
| Genetically independent CpGs (N=150,971); mean ΦST = 0.046 ± 0.001 (p < 0.001) | | | |
|  | New Caledonia | Fiji | French Polynesia |
| New Caledonia |  |  |  |
| Fiji | 0.047 |  |  |
| French Polynesia | 0.047 | 0.045 |  |

**Table S6**. Pairwise *F*_ST_ between archipelagos in *A.hyacinthus*

| *Acropora hyacinthus* (N=100; 12,613 SNPs) | | | |
| --- | --- | --- | --- |
| **Population** | **Pairwise *F*_ST_** | **CI lower** | **CI upper** |
| New Caledonia / Fiji | 0.055 | 0.053 | 0.057 |
| New Caledonia / French Polynesia | 0.078 | 0.075 | 0.080 |
| Fiji / French Polynesia | 0.071 | 0.069 | 0.074 |

**Table S7**. Heterozygosity of *A.hyacinthus* at archipelago level. Observed heterozygosity differed significantly among *A.hyacinthus* populations (Kruskal-Wallis test, χ² = 77.37, p < 0.001). Pairwise comparisons (Dunn-test with BH correction) showed significant differences between all groups.

| *Acropora hyacinthus* (N=100; 12,613 SNPs) | | | | | |
| --- | --- | --- | --- | --- | --- |
| **Population** | ***H*o** | ***H*e** | ***F*_IS_** | **CI lower** | **CI upper** |
| New Caledonia | 0.146 | 0.122 | -0.197 | -0.203 | -0.192 |
| Fiji | 0.135 | 0.114 | -0.191 | -0.197 | -0.185 |
| French Polynesia | 0.142 | 0.121 | -0.178 | -0.184 | -0.173 |

**Table S8**. Pairwise Φ_ST_ between sites of New Caledonia of *P.acuta* colonies. For each dataset (SNPs (A); genetically associated CpGs (B); genetically independent CpGs (C)), two tables are presented: the first contains the Φ_ST_ values, and the second contains the p-values associated with the comparisons. Significant comparisons are highlighted in red.

A. Pairwise Φ_ST_ on SNPs (N=21; 7,477 SNPs); mean 0.032 ± 0.03

|  | Citrons | Prony | Naia | Maa | B1 | R1 |
| --- | --- | --- | --- | --- | --- | --- |
| Citrons |  |  |  |  |  |  |
| Prony | -0,0046 |  |  |  |  |  |
| Naia | -0,0212 | -0,0262 |  |  |  |  |
| Maa | -0,0220 | -0,0253 |  |  |  |  |
| B1 | 0,0301 | 0,0207 | 0,0441 | 0,0295 |  |  |
| R1 | 0,0387 | 0,0238 | 0,1661 | 0,1595 | 0,0837 |  |

| ***p*-values** | Citrons | Prony | Naia | Maa | B1 | R1 |
| --- | --- | --- | --- | --- | --- | --- |
| Citrons |  |  |  |  |  |  |
| Naia | 0,7556 |  |  |  |  |  |
| Prony | 0,5986 | 0,8756 |  |  |  |  |
| Maa | 0,8056 | 0,88 |  |  |  |  |
| B1 | 0,0033 | 0,0017 | 0,1428 | 0,2841 |  |  |
| R1 | 0,0718 | 0,1366 | 0,3275 | 0,3408 | 0,0373 |  |

B. Pairwise Φ_ST_ on genetically associated CpGs (N=46; 47,296 CpGs); mean 0.07 ± 0.048

|  | Citrons | Naia | Prony | Maa | B1 | R1 |
| --- | --- | --- | --- | --- | --- | --- |
| Citrons |  |  |  |  |  |  |
| Naia | 0,0696 |  |  |  |  |  |
| Prony | 0,0128 | 0,0842 |  |  |  |  |
| Maa | 0,0342 | 0,0955 | 0,0399 |  |  |  |
| B1 | 0,0447 | 0,1128 | 0,0150 | 0,0596 |  |  |
| R1 | 0,0908 | 0,1852 | 0,0273 | 0,1312 | 0,0417 |  |

| ***p*-values** | Citrons | Naia | Prony | Maa | B1 | R1 |
| --- | --- | --- | --- | --- | --- | --- |
| Citrons |  |  |  |  |  |  |
| Naia | 0,0013 |  |  |  |  |  |
| Prony | 0,0774 | 2,00E-04 |  |  |  |  |
| Maa | 0,0116 | 1,00E-04 | 8,00E-04 |  |  |  |
| B1 | 0,0001 | 0,0001 | 0,0026 | 0,0001 |  |  |
| R1 | 0,0025 | 0,0019 | 0,0444 | 0,0015 | 0,0022 |  |

C. Pairwise Φ_ST_ on genetically independent CpGs (N=46; 59,928 CpGs); mean 0.05 ± 0.023

|  | Citrons | Naia | Prony | Maa | B1 | R1 |
| --- | --- | --- | --- | --- | --- | --- |
| Citrons |  |  |  |  |  |  |
| Port Naia | 0,0553 |  |  |  |  |  |
| Prony | 0,0113 | 0,0493 |  |  |  |  |
| Maa | 0,0489 | 0,0925 | 0,0386 |  |  |  |
| B1 | 0,0539 | 0,0932 | 0,0347 | 0,0623 |  |  |
| R1 | 0,0411 | 0,0718 | 0,0229 | 0,0500 | 0,0270 |  |

| ***p*-values** | Citrons | Naia | Prony | Maa | B1 | R1 |
| --- | --- | --- | --- | --- | --- | --- |
| Citrons |  |  |  |  |  |  |
| Port Naia | 3,00E-04 |  |  |  |  |  |
| Prony | 0,0709 | 6,00E-04 |  |  |  |  |
| Maa | 0,0099 | 2,00E-04 | 0,004 |  |  |  |
| B1 | 1,00E-04 | 1,00E-04 | 0,0001 | 5,00E-04 |  |  |
| R1 | 0,0033 | 0,0015 | 0,0025 | 0,0199 | 0,0193 |  |

**Table S9***.* Pairwise Φ_ST_ between sites in Fiji of *P.acuta* colonies. For each dataset (SNPs (A); genetically associated CpGs (B); genetically independent CpGs (C)), two tables are presented: the first contains the Φ_ST_ values, and the second contains the p-values associated with the comparisons. Significant comparisons are highlighted in red.

A. Pairwise Φ_ST_ on SNPs (N=36; 9,444 SNPs); mean 0.0007 ± 0.001

|  | Namada | Vatuolalai | Votua | Tagage |
| --- | --- | --- | --- | --- |
| Namada |  |  |  |  |
| Vatuolalai | 0,0010 |  |  |  |
| Votua | 0,0015 | 0,0007 |  |  |
| Tagage | -0,0003 | -0,0006 | 0,0019 |  |

| ***p*-values** | Namada | Vatuolalai | Votua | Tagage |
| --- | --- | --- | --- | --- |
| Namada |  |  |  |  |
| Vatuolalai | 0,379 |  |  |  |
| Votua | 0,3078 | 0,4458 |  |  |
| Tagage | 0,5055 | 0,5503 | 0,357 |  |

B. Pairwise Φ_ST_ on genetically associated CpGs (N=37; 43,336 CpGs); mean 0.001 ± 0.004

|  | Namada | Vatuolalai | Votua | Tagage |
| --- | --- | --- | --- | --- |
| Namada |  |  |  |  |
| Vatuolalai | -0,0049 |  |  |  |
| Votua | 0,00003 | -0,0064 |  |  |
| Tagage | 0,0059 | -0,0018 | -0,0003 |  |

| ***p*-values** | Namada | Vatuolalai | Votua | Tagage |
| --- | --- | --- | --- | --- |
| Namada |  |  |  |  |
| Vatuolalai | 0,6469 |  |  |  |
| Votua | 0,3369 | 0,6274 |  |  |
| Tagage | 0,1387 | 0,4329 | 0,3944 |  |

C. Pairwise Φ_ST_ on independent CpGs (N=37; 67,988 CpGs); mean 0.046 ± 0.005

|  | Namada | Vatuolalai | Votua | Tagage |
| --- | --- | --- | --- | --- |
| Namada |  |  |  |  |
| Vatuolalai | 0,0002 | NA |  |  |
| Votua | 0,0013 | 0,0003 |  |  |
| Tagage | 0,0121 | 0,0069 | 0,0070 |  |

| ***p*-values** | Namada | Vatuolalai | Votua | Tagage |
| --- | --- | --- | --- | --- |
| Namada |  |  |  |  |
| Vatuolalai | 0,422 |  |  |  |
| Votua | 0,2626 | 0,4076 |  |  |
| Tagage | 0,0107 | 0,1171 | 0,119 |  |

**Table S10.** Pairwise Φ_ST_ between sites in French Polynesia of *P.acuta* colonies. For each dataset (SNPs (A); genetically associated CpGs (B); genetically independent CpGs (C)), two tables are presented: the first contains the Φ_ST_ values, and the second contains the p-values associated with the comparisons. Significant comparisons are highlighted in red.

A. Pairwise Φ_ST_ on SNPs (N=24; 4,028 SNPs); 0.023 ± 0.02

|  | Pao | Jet | PK17 | Vairao | Tautira |
| --- | --- | --- | --- | --- | --- |
| Pao |  |  |  |  |  |
| Jet | 0,0256 |  |  |  |  |
| PK17 | 0,0094 | 0,0372 |  |  |  |
| Vairao | 0,0316 | 0,0486 | 0,0159 |  |  |
| Tautira | 0,0233 | 0,0412 | -0,0209 | 0,0216 |  |

| ***p*-values** | Pao | Jet | PK17 | Vairao | Tautira |
| --- | --- | --- | --- | --- | --- |
| Pao |  |  |  |  |  |
| Jet | 0,0409 |  |  |  |  |
| PK17 | 0,1081 | 0,0532 |  |  |  |
| Vairao | 0,0081 | 7,00E-04 | 0,1842 |  |  |
| Tautira | 0,0181 | 0,0373 | 0,7037 | 0,1187 |  |

B. Pairwise Φ_ST_ on genetically associated CpGs (N=44; 47,679 CpGs); mean 0.038 ± 0.015

|  | Jet | Pao | PK17 | Vairao | Tautira |
| --- | --- | --- | --- | --- | --- |
| Jet |  |  |  |  |  |
| Pao | 0,0236 |  |  |  |  |
| PK17 | 0,0391 | 0,0396 |  |  |  |
| Vairao | 0,0137 | 0,0250 | 0,0297 |  |  |
| Tautira | 0,0433 | 0,0646 | 0,0522 | 0,0458 |  |

| ***p*-values** | Jet | Pao | PK17 | Vairao | Tautira |
| --- | --- | --- | --- | --- | --- |
| Jet |  |  |  |  |  |
| Pao | 0,0049 |  |  |  |  |
| PK17 | 0,0001 | 0,0001 |  |  |  |
| Vairao | 0,0015 | 1,00E-04 | 0,0001 |  |  |
| Tautira | 7,00E-04 | 5,00E-04 | 3,00E-04 | 3,00E-04 |  |

C. Pairwise Φ_ST_ on genetically independent CpGs (N=44; 63,742 CpGs); mean 0.042 ± 0.018

|  | Jet | Pao | PK17 | Vairao | Tautira |
| --- | --- | --- | --- | --- | --- |
| Jet |  |  |  |  |  |
| Pao | 0,0303 |  |  |  |  |
| PK17 | 0,0234 | 0,0371 |  |  |  |
| Vairao | 0,0268 | 0,0339 | 0,0243 |  |  |
| Tautira | 0,0560 | 0,0747 | 0,0556 | 0,0623 |  |

| ***p*-values** | Jet | Pao | PK17 | Vairao | Tautira |
| --- | --- | --- | --- | --- | --- |
| Jet |  |  |  |  |  |
| Pao | 0,0012 |  |  |  |  |
| PK17 | 0,0066 | 0,0001 |  |  |  |
| Vairao | 0,0001 | 1,00E-04 | 8,00E-04 |  |  |
| Tautira | 0,0012 | 8,00E-04 | 0,0025 | 3,00E-04 |  |

**Table S11**. Pairwise Φ_ST_ between sites in New Caledonia of *A.hyacinthus* colonies. For each dataset (SNPs (A); genetically associated CpGs (B); genetically independent CpGs (C)), two tables are presented: the first contains the Φ_ST_ values, and the second contains the p-values associated with the comparisons. Significant comparisons are highlighted in red.

A. Pairwise Φ_ST_ on SNPs (N=29; 5,187 SNPs); mean 0.002 ± 0.002

|  | Prony | Maitre | Snark | Crouy | R2 |
| --- | --- | --- | --- | --- | --- |
| Prony |  |  |  |  |  |
| Maitre | 0,0009 |  |  |  |  |
| Snark | 0,0014 | 0,0011 |  |  |  |
| Crouy | 0,0008 | -0,0005 | 0,0059 |  |  |
| R2 | 0,0019 | 0,0018 | 0,0045 | 0,0046 |  |

| ***p*-values** | Prony | Maitre | Snark | Crouy | R2 |
| --- | --- | --- | --- | --- | --- |
| Prony |  |  |  |  |  |
| Maitre | 0,2186 |  |  |  |  |
| Snark | 0,2688 | 0,3329 |  |  |  |
| Crouy | 0,343 | 0,6104 | 0,049 |  |  |
| R2 | 0,2313 | 0,2116 | 0,1967 | 0,0915 |  |

B. Pairwise Φ_ST_ on associated CpGs (N=29; 48,562 CpGs), mean 0.01 ± 0.008

|  | Prony | Maitre | Snark | Crouy | R2 |
| --- | --- | --- | --- | --- | --- |
| Prony |  |  |  |  |  |
| Maitre | 0,0004 |  |  |  |  |
| Snark | 0,0157 | 0,0119 |  |  |  |
| Crouy | -0,0003 | 0,0001 | 0,0157 |  |  |
| R2 | 0,0106 | 0,0093 | 0,0220 | 0,0154 |  |

| ***p*-values** | Prony | Maitre | Snark | Crouy | R2 |
| --- | --- | --- | --- | --- | --- |
| Prony |  |  |  |  |  |
| Maitre | 0,2926 |  |  |  |  |
| Snark | 0,0191 | 0,0279 |  |  |  |
| Crouy | 0,5128 | 0,4323 | 0,0167 |  |  |
| R2 | 0,0429 | 0,0244 | 0,1034 | 0,0179 |  |

C. Pairwise Φ_ST_ on independent CpGs (N=29; 158,528 CpGs); mean 0.007 ± 0.004

|  | Prony | Maitre | Snark | Crouy | R2 |
| --- | --- | --- | --- | --- | --- |
| Prony |  |  |  |  |  |
| Maitre | 0,0012 |  |  |  |  |
| Snark | 0,0106 | 0,0093 |  |  |  |
| Crouy | 0,0041 | 0,0036 | 0,0110 |  |  |
| R2 | 0,0051 | 0,0104 | 0,0137 | 0,0044 |  |

| ***p*-values** | Prony | Maitre | Snark | Crouy | R2 |
| --- | --- | --- | --- | --- | --- |
| Prony |  |  |  |  |  |
| Maitre | 0,2037 |  |  |  |  |
| Snark | 0,0478 | 0,0099 |  |  |  |
| Crouy | 0,0557 | 0,0474 | 0,0338 |  |  |
| R2 | 0,1282 | 0,0041 | 0,1023 | 0,1582 |  |

**Table S12**. Pairwise Φ_ST_ between sites in Fiji of *A.hyacinthus* colonies. For each dataset (SNPs (A); genetically associated CpGs (B); genetically independent CpGs (C)), two tables are presented: the first contains the Φ_ST_ values, and the second contains the p-values associated with the comparisons. Significant comparisons are highlighted in red.

A. Pairwise Φ_ST_ on SNPs (N=31; 4,538 SNPs); mean 0.025 ± 0.02

|  | Vatuolalai | Tagage | Namada | Votua |
| --- | --- | --- | --- | --- |
| Vatuolalai |  |  |  |  |
| Tagage | 0,036 |  |  |  |
| Namada | 0,051 | 0,016 |  |  |
| Votua | 0,032 | -0,002 | 0,016 |  |

| ***p*-values** | Vatuolalai | Tagage | Namada | Votua |
| --- | --- | --- | --- | --- |
| Vatuolalai |  |  |  |  |
| Tagage | 0,0025 |  |  |  |
| Namada | 0,0945 | 0,0407 |  |  |
| Votua | 0,0136 | 0,8732 | 0,2242 |  |

B. Pairwise Φ_ST_ on associated CpGs (N=31; 44,275 CpGs); mean 0.01 ± 0.01

|  | Vatuolalai | Tagage | Namada | Votua |
| --- | --- | --- | --- | --- |
| Vatuolalai |  |  |  |  |
| Tagage | 0,016 |  |  |  |
| Namada | 0,024 | 0,003 |  |  |
| Votua | 0,016 | 0,0005 | 0,003 |  |

| ***p*-values** | Vatuolalai | Tagage | Namada | Votua |
| --- | --- | --- | --- | --- |
| Vatuolalai |  |  |  |  |
| Tagage | 0,0012 |  |  |  |
| Namada | 0,0019 | 0,1226 |  |  |
| Votua | 0,0018 | 0,328 | 0,1462 |  |

C. Pairwise Φ_ST_ on independent CpGs (N=31; 166,018 CpGs); mean 0.03 ± 0.002

|  | Vatuolalai | Tagage | Namada | Votua |
| --- | --- | --- | --- | --- |
| Vatuolalai |  |  |  |  |
| Tagage | 0,053 |  |  |  |
| Namada | 0,058 | 0,008 |  |  |
| Votua | 0,044 | 0,005 | 0,008 |  |

| ***p*-values** | Vatuolalai | Tagage | Namada | Votua |
| --- | --- | --- | --- | --- |
| Vatuolalai |  |  |  |  |
| Tagage | 8,00E-04 |  |  |  |
| Namada | 0,0024 | 0,0067 |  |  |
| Votua | 0,002 | 0,0077 | 0,0156 |  |

**Table S13**. Pairwise Φ_ST_ between sites in French Polynesia of *A.hyacinthus* colonies. For each dataset (SNPs (A); genetically associated CpGs (B); genetically independent CpGs (C)), two tables are presented: the first contains the ΦST values, and the second contains the p-values associated with the comparisons. Significant comparisons are highlighted in red.

A. Pairwise Φ_ST_ on SNPs (N=40; 7,608 SNPs); mean 0.0007 ± 0.001

|  | Jet | PK17 | Vairao | Tautira |
| --- | --- | --- | --- | --- |
| Jet |  |  |  |  |
| PK17 | 0,0019 |  |  |  |
| Vairao | -0,0004 | 0,0013 |  |  |
| Tautira | -0,0006 | 0,0017 | 0,0004 |  |

| ***p*-values** | Jet | PK17 | Vairao | Tautira |
| --- | --- | --- | --- | --- |
| Jet |  |  |  |  |
| PK17 | 0,018 |  |  |  |
| Vairao | 0,6501 | 0,1776 |  |  |
| Tautira | 0,7641 | 0,0667 | 0,3622 |  |

B. Pairwise Φ_ST_ on associated CpGs (N=40; 57,756 CpGs); mean 0.003 ± 0.002

|  | Jet | PK17 | Vairao | Tautira |
| --- | --- | --- | --- | --- |
| Jet |  |  |  |  |
| PK17 | 0,0008 |  |  |  |
| Vairao | 0,0054 | 0,0051 |  |  |
| Tautira | 0,0027 | 0,0025 | 0,0004 |  |

| ***p*-values** | Jet | PK17 | Vairao | Tautira |
| --- | --- | --- | --- | --- |
| Jet |  |  |  |  |
| PK17 | 0,2133 |  |  |  |
| Vairao | 0,0092 | 0,0016 |  |  |
| Tautira | 0,0084 | 0,0569 | 0,3324 |  |

C. Pairwise Φ_ST_ on independent CpGs (N=40; 156,922 CpGs); mean 0.0074 ± 0.002

|  | Jet | PK17 | Vairao | Tautira |
| --- | --- | --- | --- | --- |
| Jet |  |  |  |  |
| PK17 | 0,0036 |  |  |  |
| Vairao | 0,0075 | 0,0080 |  |  |
| Tautira | 0,0073 | 0,0100 | 0,0080 |  |

| ***p*-values** | Jet | PK17 | Vairao | Tautira |
| --- | --- | --- | --- | --- |
| Jet |  |  |  |  |
| PK17 | 0,0449 |  |  |  |
| Vairao | 9,00E-04 | 0,0279 |  |  |
| Tautira | 0,0011 | 0,016 | 0,0171 |  |

**Data S1**. **(Separate file)**. Metadata of *P.acuta* and *A.hyacinthus* (sample_id, archipelago and site of origin). For *P.acuta*, the table contains the clones identified following the relatedness analysis in column E.
